# ChIP-Hub: an Integrative Platform for Exploring Plant Regulome

**DOI:** 10.1101/768903

**Authors:** Dijun Chen, Liang-Yu Fu, Peijing Zhang, Ming Chen, Kerstin Kaufmann

**Affiliations:** Department for Plant Cell and Molecular Biology, Institute for Biology, Humboldt-Universität zu Berlin, 10115 Berlin, Germany; School of Life Sciences, Nanjing University, Nanjing 210023, China; Department of Bioinformatics, College of Life Sciences, Zhejiang University, Hangzhou 310058, China

## Abstract

Plant genomes encode a complex and evolutionary diverse regulatory grammar that forms the basis for most life on earth. A wealth of regulome and epigenome data have been generated in various plant species, but no common, standardized resource is available so far for biologists. Here we present ChIP-Hub, an integrative web-based platform in the ENCODE standards that bundles publicly available datasets reanalyzed from >40 plant species, allowing visualization and meta-analysis.

## Main

Genome-wide charting of transcription factor binding and epigenetic status has become widely used to study gene-regulatory programs in animals and plants. Accordingly, a tremendous amount of data have been generated by several large consortiums (such as the ENCODE consortium in human^1^ and mouse^2^, as well as the modENCODE consortium in fly^3^ and nematode^4^) or various smaller projects (such as the fruitENCODE project in flowering plants^5^). Several databases^6–8^ were recently established for visualization and efficient deployment of public ChIP-seq data by the research community. However, no comprehensive resource is available for plant research. Another major bottleneck in current plant research is the lack of a standardized routine for evaluation and analysis of ChIP-seq data. Therefore, the comparison of data generated by different laboratories is not straightforward, hampering data integration to generate novel hypotheses for further investigation.

Here, we present an integrative web-based platform (ChIP-Hub, http://www.chip-hub.org) for exploring the comprehensive reanalysis of >5600 individual datasets in more than 40 plant species, including data to study both transcription-factor (TF) binding and histone modifications (**Fig. 1a** and **Supplementary Fig. 1**). To this end, we adapted the working standards provided by the ENCODE consortium^9^ to set up computational pipelines (**Fig. 1b**) and to systematically reanalyze public ChIP-seq and DAP-seq^10,11^ experiments in plants, with careful manual curation through assessing ∼360 original publications (**Fig. 1a,b**). We systematically evaluate the data quality of individual experiments (n=3078; **Fig. 1c**). Although 93.3% of the experiments have been published in peer-reviewed journals, nearly 25% of the experiments lack control datasets, 36.7% have a low sequencing depth that limits peak calling, and only 37.5% have replicates. Problems of low sequencing depth and lack of controls or replicates is more obvious in the earlier studies (**Supplementary Fig. 2**). Nevertheless, most of the evaluated experiments readily meet a variety of quality specifications based on the ENCODE criteria^9^ (**Fig. 1c**).

**Figure 1.**
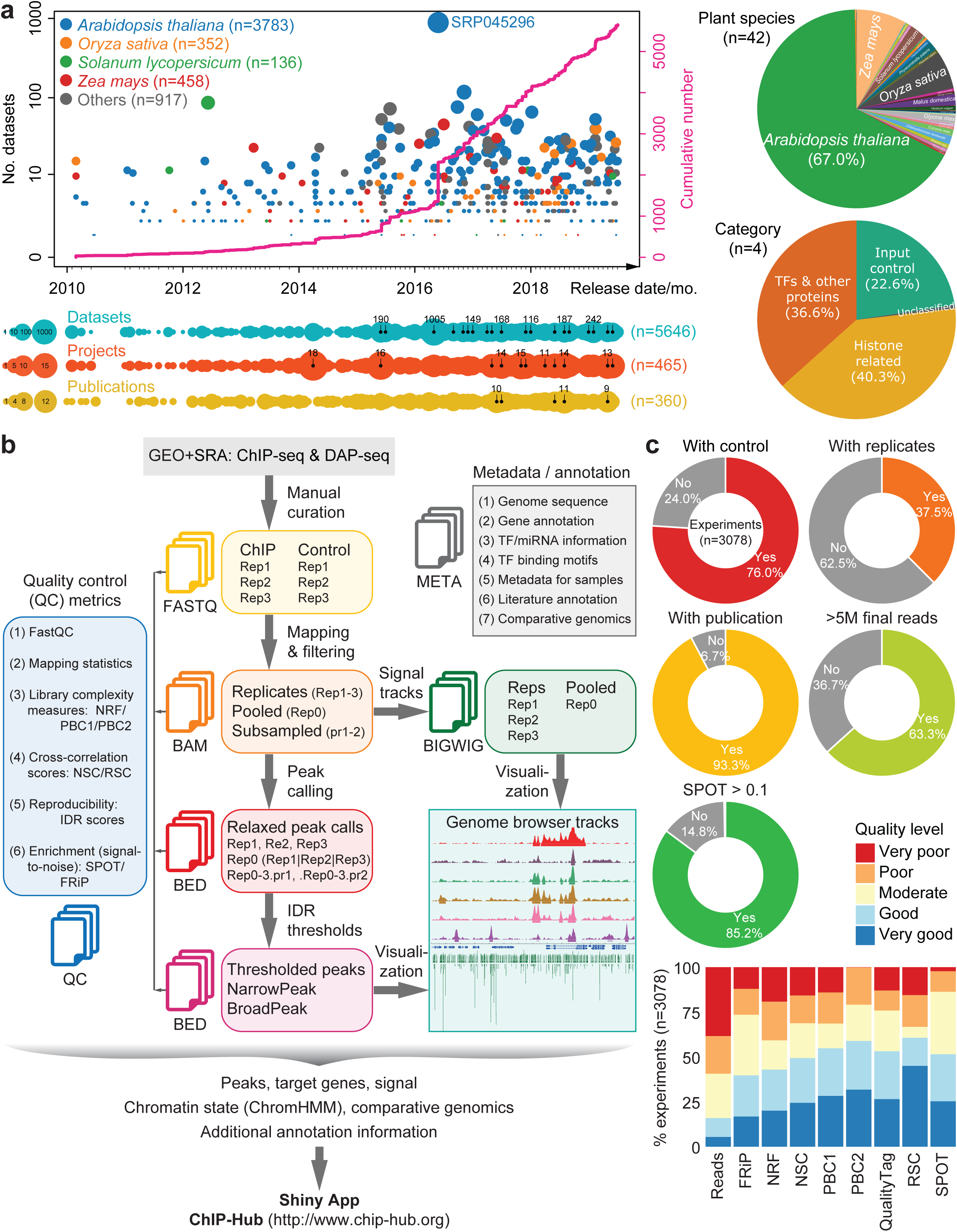
Reanalysis and evaluation of published regulome and epigenome experiments in plants. (**a**) Overview of published regulome and epigenome data in plants. The scatter plot (top left) shows the number of datasets over time (in month [mo.]), as colored by the top representative plant species. Each data point represents one SRA BioProject. The cumulative number is also shown. An overview of the number of datasets, publications and BioProjects over time is shown in the timeline plots (bottom left). The right pie charts showing the distribution of datasets by plant species (top) or by sample categories (bottom). (**b**) A standardized, semi-automatic analysis pipeline developed for regulome and epigenome experiments. We adapted the working standards provided by the ENCODE consortium^9^ to set up the computational pipeline, including read mapping, peak calling and subsequent statistical treatment of replicates. The resulting data are further integrated by ChromHMM^14^ for each plant species. All the metadata as well as analyzed data are bundled in our Shiny application ChIP-Hub for visualization and meta-analysis. (**c**) Evaluation of published regulome and epigenome experiments (n=3087) in plants. Donut charts (top) show different aspects of evaluation of the published experiments. The bottom bar chart shows the quality of experiments based on various quality metrics proposed by the ENCODE consortium^9^. Refer to **Supplementary Fig. 18** for the definition of metrics categories. Note that all the numbers in this figure were summarized based on data collected on July 9th, 2019. SRA: sequence read archive; SPOT: signal portion of tags; FRiP: fraction of reads in peaks; NSC: normalized strand cross-correlation coefficient. RSC: relative Strand cross-correlation coefficient; NRF: non-redundant fraction; PBC1/2: PCR bottlenecking coefficients 1/2.

We identified a total of 23.7 million high-confidence peaks (with an IDR, Irreproducible Discovery Rate^12^, < 0.05; see **Methods**) from experiments for annotated TFs or widely-investigated histone H3 modifications. For genomes with more than 15 distinct experiments, the number of identified TF binding events or histone-modified genomic regions varies from 0.21 million (*Chlamydomonas reinhardtii*; experiments n=32) to 8.5 million (*Arabidopsis thaliana*; n=1647); the fraction of genome associated with potential functional elements shows an average of 18.0 % (**Fig. 2a**), with comparable proportions found in the mouse (12.6%) and human (∼20%) genomes^1,2^. However, the proportion may be far underestimated for most plant genomes since many regulators have not yet been investigated. Of note, about 1600 individual experiments for >640 different factors and 15 histone modifications have been generated in *Arabidopsis* (**Fig. 2b**), resulting in functionally annotated genomic regions encompassing at least 82.0% of the *Arabidopsis* genomic sequence in aggregate (**Fig. 2a**). Interestingly, 58.1% of *Arabidopsis* genome is annotated as potential regulatory regions based on 291 ChIP-seq experiments for 129 distinct TFs (**Supplementary Fig. 3** and **Supplementary Table 1**), suggesting pervasive genome regulation in *Arabidopsis*. Integrative analysis of TF-bound genomic regions and target genes revealed potential TF co-associations (**Supplementary Figs. 4** and **5**) and comprehensive miRNA-mediated feed-forward loops^13^ (FFLs; **Supplementary Fig. 6**).

**Figure 2.**
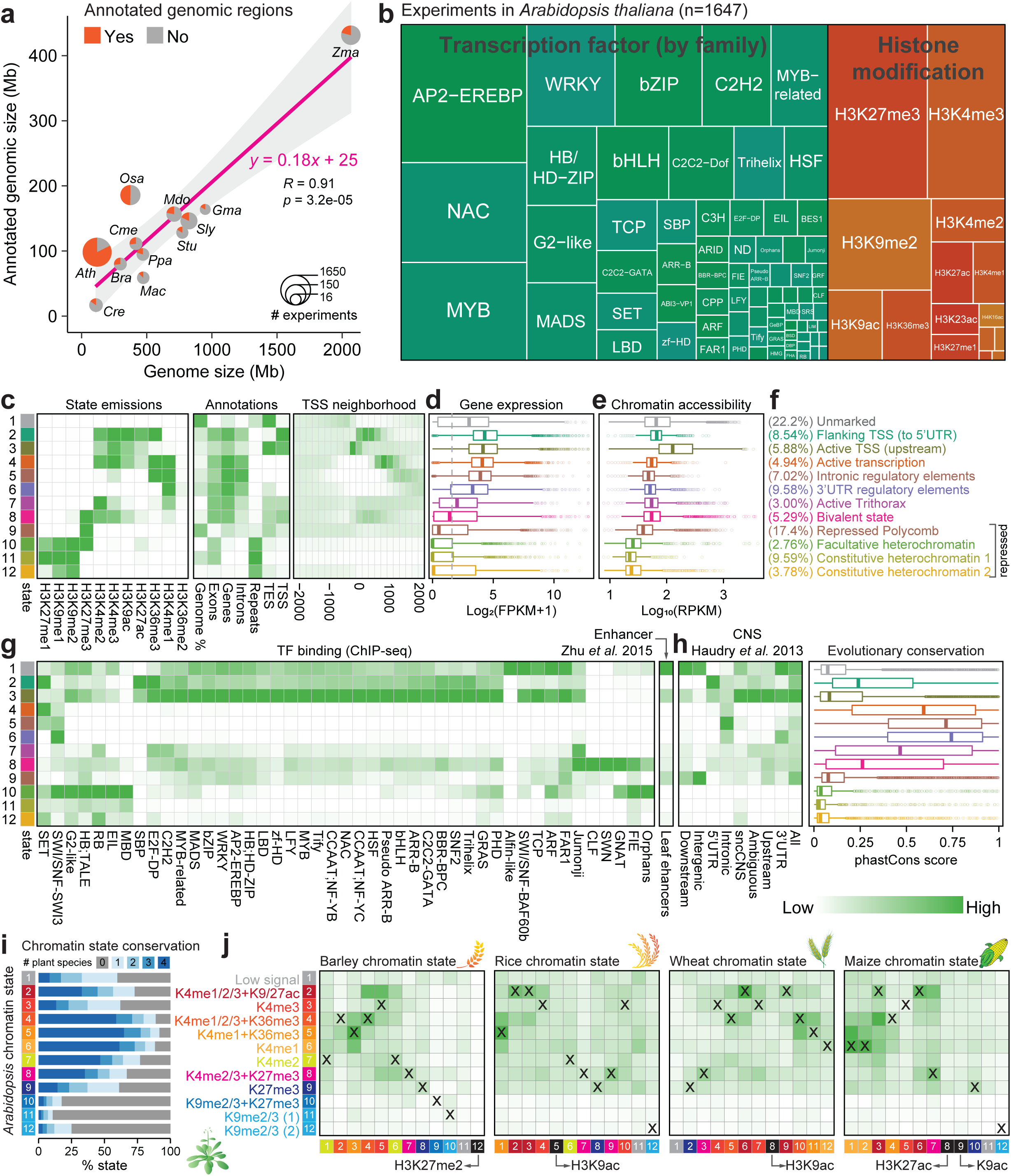
Systematic annotation of regulome and epigenome experiments. (**a**) Annotated genomic regions versus the genome size. Pie charts show the percentage of genomes annotated by ChIP-seq data. Only genomes with >15 experiments are shown. Full names of genomes can be found in **Supplementary Table 4**. (**b**) Treemap showing the classification of experiments in *Arabidopsis thaliana* according to the types of histone modifications or transcription factor (TF) families. (**c-h**) Definition and enrichment for a 12-state ChromHMM^14^ model based on eleven histone modification marks in *Arabidopsis* vegetative-related tissues. Darker green color in the heatmaps indicates a higher probability or enrichment. In the plots, each row corresponds to a different state (in different colors), and each column corresponds to a different mark, a genomic annotation (**c**), gene expression patterns (**d**), chromatin accessibility^19^(**e**), TF binding for a different TF families and leaf enhancers^20^(**g**), or conservation information (**h**). Percentage and description of states summarized based on the overall enrichment of different categories of annotations (see **Supplementary Text** for discussion) are shown in (**f**). Gene expression data from ref.^20^; conserved noncoding sequences (CNSs) and phastCons conservation score (based on nine-way multiple alignment) between *Arabidopsis* and other crucifers from ref.^21^. (**i,j**) Chromatin state conservation between *Arabidopsis* and other four plant species with annotated states, in vegetative-related tissues. (**i**) Bar chart showing the percentage of conserved *Arabidopsis* chromatin states. The number of conserved plants is distinctly colored. Colors for states are explained in (**j**). (**j**) Enrichment of chromatin state conservation between *Arabidopsis* (row) and other species (column). Pairwise enrichment score was calculated based on Jaccard statistics, which measures the ratio of the number of conserved base pairs to the number of base pairs in union. Darker green in the heatmap indicates a higher enrichment. States with similar compositions of histone modification marks are colored in the same way among different plant species. Matched states between *Arabidopsis* and other species are labeled as “X”. Chromatin states without matched states in *Arabidopsis* are indicated in black. Unmarked states are colored in grey. Annotation of chromatin states in barley, rice, wheat and maize can be found in **Supplementary Figs. 9-13**.

In order to predict the functional relevance of the genomic regions marked by histone modifications, we generated integrated maps of chromatin states in vegetative-, reproductive- or root-related tissues of wide-type plants for genomes with at least five distinct marks (**Supplementary Table 3** and **Supplementary Fig. 7**), using ChromHMM^14^ to segment the genome into distinct combination of histone modification marks (**Supplementary Fig. 8**). As a proof of concept, a 12-state model was trained in *Arabidopsis* vegetative-related tissues (**Fig. 2c**). The resulting “marked” states included six active states, four repressed states and a bivalent state that showed distinct levels of TF binding and enrichment for evolutionary conserved noncoding sequences (**Fig. 2d-h**; see also **Supplementary Text**), accounting for 77.8% of the genome (**Fig. 2f**) and covering all the major states identified in previous studies^15–17^. The generation of these tissue-specific maps of chromatin states (**Fig. 2c-h** and **Supplementary Figs. 9-13**) also offers an unprecedented level of comparison of genomic features among different plant species. We thus tracked the evolution of chromatin states in vegetative-related tissues across five plant species (i.e., *Arabidopsis*, rice, barley, wheat and maize) using *Arabidopsis* as a reference (see **Methods**). We observed that most *Arabidopsis* chromatin states (excepted heterochromatin-related states) were highly conserved in other plant species (**Fig. 2i**). For example, orthologous sequences were found for 61.1% of Polycomb-repressed regions in at least one of the compared species. Moreover, we found significant epigenomic conservation at orthologous chromatin state-marked regions (**Fig. 2j**), consistent with results in human^18^.

To make our data easily accessible to external users, we have developed an integrated Shiny application to explore all the reanalyzed data sets. Additional data (e.g., sample metadata, references, TF genes, miRNAs, TF motifs, chromatin states and comparative genomics) were also collected and deposited in the database (**Fig. 1b**). Therefore, the resources are bundled in a well-accessible application that also allows visualization and meta-analyses (**Supplementary Figs. 14-17**).

In conclusion, ChIP-Hub serves as a comprehensive data portal to explore plant regulomes. A routine to maintain and update ChIP-Hub in the future has been established. We hope that ChIP-Hub will not only allow experimental biologists from various fields to comprehensively use all available regulome and epigenome information to get novel insights into their specific questions, but also allow theoretical biologists to model regulatory relationships under a specific conditions and developmental states.

## Acknowledgements

The authors acknowledge the North-German Supercomputing Alliance (HLRN) and the Center for Information Technology and Media Management (ZIM) at Potsdam University for providing high performance computing (HPC) resources that have contributed to the research results reported in this paper. We would like to thank all the data contributors who make this project possible. We would like to thank all the group members in Kaufmann lab for helpful discussion and suggestions. Kerstin Kaufmann wishes to thank the Alexander-von-Humboldt foundation and the Federal Ministry of Education and Research for support.

## Author contributions

D.C. and K.K. conceived and designed the study. D.C., L.-Y.F. and P.Z. annotated the sample metadata. D.C. implemented the computational analysis pipeline and developed the Shiny application with contribution from M.C.. D.C. and L.-Y.F. performed the analyses. D.C. and K.K. wrote the manuscript. All authors reviewed and approved the submitted version.

## Methods

### Data source, curation and collection

Metadata of ChIP-seq and DAP-seq samples (equivalent to datasets, accession numbers start with SRX/ERX/DRX) and projects (start with SRP/ERP/DRP) were retrieved from NCBI SRA (https://www.ncbi.nlm.nih.gov/sra), BioSample (https://www.ncbi.nlm.nih.gov/biosample), BioProject (https://www.ncbi.nlm.nih.gov/bioproject) and/or GEO (https://www.ncbi.nlm.nih.gov/geo) databases. ChIP-Hub has a focus on data in “green plants” (i.e., only considering plants in the taxonomy tree with a root ID 33090). Only data generated by Illumina platforms were kept. Firstly, each dataset was associated with publication(s) if available (more than 90% samples can be linked with publications). Then, each dataset was manually curated to determine its investigated factor (i.e., which TF or histone modification mark), its experimental type (whether ChIP or control) and its associated replicates (experiment may have several replicates), based on the metadata and the original publications. Note that it is important to manually check the metadata based on its corresponding publication since some metadata was misannotated in the database. For example, the dataset SRX4063234 in fact contains two different samples, one for ChIP experiment (SRR7142417) and another for control experiment (SRR7142416). In this case, “Run” accessions (start with SRR/ERR/DRR) were instead used as sample accessions (ca. 250 of such cases). For datasets without related publications so far, they were marked as a “unconfirmed” status and would be regularly checked in the future. In general, one experiment may contain replicate samples (i.e., datasets), ChIP sample(s) as well as input control sample(s) and it was designed to investigate regulation of a specific factor (e.g., TF or histone modification) of interest under specific conditions. In the analysis (see the section below), each experiment was processed independently. Furthermore, annotation information for investigated factors was also manually curated. Broadly, factors are grouped into “TFs and other proteins”, “histone-related” or “unclassified”. For TFs, their gene IDs and family information were also determined if applicable. Finally, a meta file was obtained for each experiment after curation (see **Supplementary Fig. 19** for examples), which is served as an input file for the ChIP-seq computation pipeline (see below).

Raw fastq files for each experiment were downloaded from the European Nucleotide Archive (ENA, https://www.ebi.ac.uk/ena) database. If fastq files were not available at ENA, raw data in the SRA format were downloaded from the SRA database and converted into fastq format using the “fastq-dump” command provided by the SRA Toolkit (version 2.5.1). The “--split-files” option was used for paired-end reads. Fastq files were further checked for completeness before submitted to analysis.

Genome sequences and gene annotations were downloaded from public databases (**Supplementary Table 4**). Additional annotation data were also included in the ChIP-Hub database in order to better annotate the regulatory factors and their regulatory networks. Annotation for miRNA genes were obtained from miRBase^22^ and their genomic locations were updated (by BLAST) based on current reference genomes. TF family information was retrieved from PlantTFDB^23^. TF DNA binding motifs were downloaded from the JASPAR^24^, CIS-BP^25^ and PlantTFDB^23^ databases and were scanned for occurrences in the genome using FIMO^26^. These data were provided as separated data tracks in the genome browser.

### ChIP-seq data processing

We followed the ChIP-seq data analysis guidelines^9^ recommended by the ENCODE project to develop computational pipeline for ChIP-seq and DAP-seq data analysis (**Fig. 1b**). The analysis pipeline consists of quality control, read mapping, peak calling and assessment of reproducibility among biological replicates and was used to analyze all annotated experiments a standardized and uniform manner. Specifically, potential adapter sequences were removed from the sequencing reads using the Trim Galore program (version 0.4.1) and the quality of sequencing data was then evaluated by FastQC (http://www.bioinformatics.babraham.ac.uk/projects/fastqc/). Clean reads were mapped to the corresponding reference genomes using Bowtie2 (version 2.2.6; ref.^27^) with parameters “-q --no-unal -- threads 8 --sensitive”. The parameter “-k” was set to 1, 2 and 3 for diploid genomes (e.g., *Oryza sativa*), tetraploid genomes (e.g., *Gossypium barbadense*) and hexaploidy genomes (e.g., *Triticum aestivum*), respectively. Redundant reads and PCR duplicates were removed using Picard tools (v2.60; http://broadinstitute.github.io/picard/) and SAMtools^28^ (version 0.1.19). Peak calling was performed using MACS2 (version 2.1.0; ref.^29^). Duplicated reads were not considered (“-

-keep-dup=1”) during peak calling in order to achieve a better specificity^30^. The shifting size (“--shift”) used in the model was determined by the analysis of cross-correlation scores using the phantompeakqualtools package (https://code.google.com/p/phantompeakqualtools/). The parameter “--call-summits” was used to call narrow peaks. For broad marks of histone modifications (including H3K36me3, H3K20me1, H3K4me1, H3K79me2, H3K79me3, H3K27me3, H3K9me3 and H3K9me1), broad peaks were also called by turning on the “--broad” parameter in MACS2. A relaxed threshold of p-value (p-value < 1e-2) was used in order to enable the correct computation of IDR (irreproducible discovery rate) values^9^, because IDR requires input peak data across the entire spectrum of high confidence (signal) and low confidence (noise) so that a bivariate model can be fitted to separate signal from noise^12^. Following the recommendations for the analysis of self-consistency and reproducibility between replicates^12^, replicate control samples (if available) were combined into one single control in the same experiment. Peak calling was applied to all replicates, pooled data (pooled replicates), pseudo-replicates (half subsample of reads) of each replicate and the pseudo-replicates of pooled sample using the same merged control as input (if applicable). By default, “reproducible” peaks across pseudo-replicates and true replicates with an IDR < 0.05 were recommend for analysis. Besides, peaks with different statistical thresholds are available upon request. For example, “significant” peaks were defined as a fold-change (fold enrichment above background) > 2 and a -log10 (q-value) > 3; while “lenient” peaks as a fold-change > 2 and a -log10 (q-value) > 2. “Relaxed” peaks without additional thresholding were also provided so that any custom threshold can be applied. All peak-based analyses in the pipeline (including peak overlapping, merging and summary) were performed using BEDTools (v2.25.0; ref.^31^).

Various metric scores were calculated to assess different aspects of the quality of experiments (https://genome.ucsc.edu/ENCODE/qualityMetrics.html and https://www.encodeproject.org/data-standards/terms/; **Fig. 1c** and **Supplementary Fig. 19**). For example, library complexity is measured using the non-redundant fraction (NRF) and PCR bottlenecking coefficients 1 and 2 (PBC1 and PBC2). The SPOT (signal portion of tags) score, characterizing the enrichment of signal for each experiment, was calculated by the Hotspot^32^ algorithm by subsampling ten million reads. Fraction of reads in peaks (FRiP), another measure of enrichment, is highly correlated with the SPOT score (**Supplementary Fig. 20**). NSC and RSC (normalized and relative strand cross-correlation coefficient) are related measures of enrichment without dependence on pre-defined peaks, which were calculated by the phantompeakqualtools program.

For visualization purpose, wiggle tracks (using pooled data across replicates) were generated by DeepTools^33^ with the “bamCoverage” program; different normalization methods (including RPKM [reads per kilobase per million mapped reads], CPM [counts per million mapped reads], BPM [bins per million mapped reads], RPGC [reads per genomic content normalized to 1x sequencing depth] and None) were used to generate different types of signal files. ChIP-seq tracks were visualized in the WashU Epigenome Browser^34^.

### Assignment of target genes

Regulatory elements (in layman’s terms, called “peaks”) were assigned to putative target genes based on the following rules. For a regulatory region overlapping with any gene(s) (protein-coding genes or miRNAs), the overlapping gene(s) were considered as its targets. Otherwise, the regulatory element was assigned to its nearest annotated gene within up to N bp, where N is the median size of intergenic regions (N was set to 3000 if the median size exceeded 3000). The start of genes (i.e., the transcription start site [TSS] of protein-coding genes and the 5’ end of miRNA precursors [pre-miRNAs]) was used to calculate the distance. In general, this approach associates a single regulatory element with no more than two genes, with a few exceptions in the case of the regulatory element overlapping multiple genes. This procedure was performed in each species independently.

### Chromatin state analysis

In order to use the collected histone modification ChIP-seq data from diverse studies for chromatin state analysis and to make the data more comparative among different plant species, only well-characterized H3-related histone modification marks (including H3K9ac, H3K27ac, H3K4me1/2/3, H3K9me1/2/3, H3K27me1/2/3 and H3K36me1/2/3) were considered and only data generated in wild-type plants were used. Furthermore, the datasets were broadly categorized into vegetative-, reproductive- and root-related samples based on their tissue specificity (**Supplementary Table 3**). In general, these broadly defined “tissue” types (termed reference tissue types) are more comparative among different plant species and difference in tissue collection by different studies can be eliminated. Although the analysis is cell type agnostic, it is informative even when the relevant cell or tissue type has not been experimentally profiled (this is the most case in plants so far). In addition, we filtered out experiments with less than 1000 called peaks and only considered plant species with at least five distinct types of histone modification marks. In summary, 251 experiments from five plant species were retained for chromatin state analysis (**Supplementary Table 3** and **Supplementary Fig. 7**).

ChromHMM^14^ (version 1.19) was applied on the ChIP-seq data of histone modifications in three reference tissue types in five plant species to learn a multivariate HMM model for segmentation of genome in each tissue type. Specifically, the called peaks were first pooled from different ChIP-seq experiments for each type of the histone modifications in each tissue type for each genome separately. Peaks within blacklist regions were excluded from the analysis. The remaining pooled peaks were then processed by the “BinarizeBed” command (with the parameter “-peaks”) into binarized data in every 200 bp window over the entire genome. Models were trained independently for each reference tissue type in each genome since the composition of marks varied in different tissue types. We ran the “LearnModel” command with the number of states ranging from 2 states to 15 states and selected an “optimal-state” model based on a rule that the number of states appeared most parsimonious in terms of clearly distinct emission properties and clear interpretability of distinction between states (**Supplementary Figs. 8-13**). Furthermore, the resulting chromatin states were interpreted based on enrichment analysis of various types of functional annotations, such as gene elements, neighboring gene expression pattern, TF binding, chromatin accessibility and predicted enhancers^20,35^. To this end, the “OverlapEnrichment” and “NeighborhoodEnrichment” commands were used in the analysis. The meaningful mnemonics of states for *Arabidopsis* vegetative-related tissues was given in **Fig. 2f**.

### Comparative genomics and cross-species comparisons

Whole-genome alignments were performed in a similar way as described in ref.^21^. Briefly, soft masked genomes were aligned to each other using the LastZ alignment algorithm^36^. Collinear alignment blocks separated by gaps of <100 kb were then “chained” according to their locations in both genomes and “netted” to choose the best sub-chain for the reference species^37^. For polyploid plants, each sub-genome was individually analyzed such that each contained non-overlapping chaining. The whole-genome alignments can be visualized together with epigenomic tracks through the integrated Epigenome Browser (see below).

Pairwise comparisons of chromatin states were performed by one-to-one mapping annotated regions between species based on the above whole-genome alignments. For regions mapped to multiple orthologous locations in the other genome (i.e., regions split over multiple alignment blocks), only the largest orthologous region in the same alignment block was considered. Marked regions were considered as conserved between species when their orthologous location in the second species overlapped a marked region by a minimum of 50%. Note that the minimum required overlap had little influence on the overall results given that the median value of overlaps is 100% and the mean value is 89.9%. To make state interpretations more comparable across different species (chromatin marks available for state prediction were slightly different among species, see **Supplementary Figs. 9-13**), the learned chromatin states were re-interpreted (**Fig. 2i,j**) based on a common set of marks as possible (**Supplementary Fig. 21**).

### Analysis of gene regulatory networks

To study gene regulatory networks (GRNs) controlled by TFs with available ChIP-seq data, we focused on a specific network motif, TF-miRNA-TF feed-forward loops (FFLs), which involves targeting of a TF to both miRNAs and miRNA target TFs. Such trifurcate regulatory circuits are of importance for fine tuning of downstream gene expression^38,39^. We highlighted the analysis on *Arabidopsis* data since a comprehensive list of TFs have been investigated by ChIP-seq experiments in this plant species (**Fig. 2c**). The methodology, however, can easily be applied to data from any other plants when more and more data are generated. In the miRNA-mediated FFLs, target genes of miRNAs were predicted by the TargetFinder tool^40^, with a prediction score cut-off value set to 4. Other relationships (i.e., TF-miRNA and TF-TF) were supported by ChIP-seq data. The final meta-network consisted of regulatory relationships among 97 master TFs, 125 miRNAs and 447 common target TFs (**Supplementary Fig. 6a** and **Supplementary Table 2**), covering nearly two-thirds of the predicted FFLs involved in flower development^13^ (**Supplementary Fig. 6b,c**).

### ChIP-Hub Shiny application

In order to efficiently use our reanalyzed data by external users, we developed an integrative web-based application (ChIP-Hub) with the Shiny framework (http://shiny.rstudio.com/), which combines the computational power of R with friendly and interactive web interfaces (**Supplementary Fig. 14**). All the sample metadata, curated metadata and analyzed result data were loaded into a MySQL database, allowing for interactive retrieval through the ChIP-Hub interface. These data were presented in tabular and chart forms in our Shiny web application. Furthermore, the data can be searched by keyword or gene to select datasets of interests. The associated result files, such as wiggle signal files, peak files and additional annotation files, can be loaded into the integrated Epigenome Browser (http://www.epiplant.hu-berlin.de/browser/) for visualization.

### Online access and updates

To make this project easier to maintain for a long life and to update in time, we have developed a semi-automatic computational program (ChIPer) for this purpose. The program regularly (in very month according to our current plan) checks whether any new datasets available in public databases. If so, the new datasets will be sent for curation via email and the curated datasets will be automatically analyzed by the data processing pipeline. New result files will be uploaded to our web server when the analysis is done. Besides, we would include more functionalities in our Shiny application as required.

### Statistics and reproducibility

If not specified, all statistical analyses and data visualization were done in R (version 3.4.1). R packages such as ggplot2 and plotly were heavily used for graphics. All the sources data for each figure can be found in the supplementary tables and the latest data can be found in our ChIP-Hub website.

### Data availability

The data can be viewed, mined and downloaded through the ChIP-Hub website (http://www.chip-hub.org).

